# Effect of Light and Water Agitation on Hatching Processes in Clown Anemonefish *Amphiprion ocellaris*

**DOI:** 10.1101/2024.02.14.580270

**Authors:** Sakuto Yamanaka, Mari Kawaguchi, Shigeki Yasumasu, Kenji Sato, Masato Kinoshita

## Abstract

Clown anemonefish (*Amphiprion ocellaris*) exhibit an environmentally cued hatching strategy, wherein parents generate water flow to encourage embryos to hatch after sunset on the eighth day post spawning. Although previous studies have shown that hatching requires complete darkness and water agitation, the regulatory mechanisms underlying these environmental cues remain elusive. This study investigated the expression patterns of hatching enzymes and how darkness and water agitation affect their secretion and the digestion of the egg envelope (chorion) in clown anemonefish. We revealed that hatching enzymes were expressed from the early developmental stages and decreased towards hatching day, similar to other fish species. The hatching gland cells were located on the tail, which may be associated with egg shape and chorion breakage. On the day of hatching, chorion digestion occurred exclusively in complete darkness, regardless of water agitation. In addition, ionomycin, a Ca^2+^ ionophore and an inducer of hatching enzyme secretion, triggered chorion digestion even under light-on conditions. Furthermore, water agitation expedited chorion digestion in the dark. In conclusion, our findings indicate that darkness triggers the secretion of hatching enzymes mediated by changes in Ca^2+^ levels, whereas water agitation assists chorion digestion activity and/or hatching enzyme secretion. These responses to environmental cues contribute to the rapid and synchronised hatching in clown anemonefish.

**Summary statement:** Light conditions and water agitation regulate the secretion of hatching enzymes and the chorion digestion activity, thereby facilitating a rapid hatching response to environmental changes in clown anemonefish.

## Introduction

Clown anemonefish (*Amphiprion ocellaris*) is a widely recognised tropical marine fish, distinguished by its vibrant orange body with three vertical white stripes. They spawn elliptical and adhesive eggs on the surfaces of hard rocks or corals in close proximity to their host sea anemone. Parents care for eggs until hatching, which includes fanning the egg clutch to ensure a fresh water supply and mouthing the eggs to remove any deceased ones (Green and McCormick, 2005). The time required for hatching depends on the temperature; at 28℃, it takes exactly eight days (Yamanaka et al., 2021). After sunset on the hatching day, parents intensify water flow towards the eggs, encouraging embryo hatching (Beldade et al., 2023). Embryos typically hatch within a few hours of sunset or lights being turned off in an aquarium (Tina et al., 2023). Previous studies have indicated that hatching requires both complete darkness and mechanical water agitation (Fobert et al., 2021, 2019; Yamanaka et al., 2021). In this study, we focused on the relationship between environmental cues and hatching processes in clown anemonefish.

The hatching process typically involves three key steps: secretion of hatching enzymes, digestion of the inner layer of the egg envelope (chorion), and escape movements from the egg capsule. Extensive research has been conducted on the enzymatic nature of hatching enzymes (Kawaguchi et al., 2013; Yasumasu et al., 1989a, 1989b), mechanisms underlying chorion digestion (Kawaguchi et al., 2010b; Yasumasu et al., 2010), and gene structure of hatching enzymes (Kawaguchi et al., 2010a; Olivotto et al., 2004; Sano et al., 2014; Yasumasu et al., 1992a, 1992b). In euteleostei, hatching enzyme genes are classified into two clades: high choriolytic enzyme (HCE) and low choriolytic enzyme (LCE) (Kawaguchi et al., 2010b). These two enzyme types cooperatively digest the inner layer of the egg envelope within half an hour to a few hours (Kawaguchi et al., 2013; Yamagami, 1988; Yasumasu et al., 1989b). Chorion digestion by hatching enzymes is a critical biochemical process for hatching in many fish species, except viviparous fish (black rockfish (Kawaguchi et al., 2008) and platyfish (Kawaguchi et al., 2015)). These biochemical processes are belived to occur just before hatching, influences the timing of hatching, and significantly affects larval dispersion and post-hatching fitness (Warkentin, 2011a, 2011b). Some fish species exhibit synchronised hatching strategies in response to environmental cues, a phenomenon known as environmentally cued hatching (ECH). For instance, California grunion *(Leuresthes tenuis*) spawns on sandy beaches in the high intertidal zone during spring tides, and the larvae employ a dramatic hatching strategy that is immediately triggered by waves during the subsequent highest tide (Griem & Martin 2000; Martin et al. 2011; Smyder & Martin 2002). Environmental factors including oxygen levels (Dimichele and Taylor, 1981, 1980), mechanical agitation (Griem & Martin 2000), and light conditions (Helvik and Walther, 1992; Yamagami, 1988) are known as cues for hatching in several fish species. However, the mechanisms by which environmental cues regulate hatching timing and the associated physiological processes remain poorly understood.

This study focused on clown anemonefish as a new model for studying ECH with the aim of elucidating how environmental cues regulate hatching timing. In clown anemonefish, although nearly all 8 dpf (day post fertilisation) embryos hatch within 30∼60 minutes of shading and shaking (Yamanaka et al., 2021), the specific hatching processes induced by environmental cues remain unclear. Considering this duration from receiving cues to hatching in clown anemonefish, we hypothesised that these environmental cues trigger the secretion of hatching enzymes and initiate the chorion digestion in clown anemonefish. We first confirmed whether clown anemonefish uses hatching enzymes to digest the egg envelope, similar to many other fish species. Second, we explored the effects of darkness and water agitation on the secretion of hatching enzymes and the digestion of the egg envelope.

## Materials and Methods

### Ethics Statement

Animal experiments were conducted in accordance with Kyoto University’s Regulations on Animal Experimentation. The care and use of fish were approved by the Kyoto University Animal Ethics Committee (Approval Code: R4–45 and R5-45).

### Fish Husbandry

The parent fish of clown anemonefish (*A.ocellaris*) were maintained in a recirculating aquarium at 27°C under a 14/10 LD cycle with lights on at 6:00 AM. They were fed a homemade pellet diet twice daily.

### Phylogenetic Analysis

We searched for homologous genes encoding hatching enzymes in the clown anemonefish genome (Genome Assembly ASM2253959v1, (Ryu et al., 2022)). Multiple amino acid sequences were aligned using MAFFT program (Katoh and Standley, 2013) and trimmed using trimAl (Capella-Gutiérrez et al., 2009). Phylogenetic trees were constructed using the maximum likelihood method with the WAG (Whelan and Goldman, 2001) + Gamma model and bootstrap analysis (1,000 replicates) under the PROTGAMMAAUTO option to determine the best-scoring topology using RAxML (Stamatakis, 2014).

### RT-qPCR

We examined the temporal expression patterns of several hatching enzyme genes, including AoLCE (XM_023278117), AoHCE1 (XM_023262355), AoHCE2 (XM_055012028), and AoHCE3 (XM_055012305) using RT-qPCR. The housekeeping gene beta-actin2 (actb2, XM_023294531) was used as an internal control for the 2^-ΔΔCT^ method. Specific primers were designed for the 3’ UTR region of each gene because of the similarity of the coding region of the hatching enzyme genes. RNA was extracted from various embryonic and larval stages (1–8 dpf embryos and 9–10 dpf larvae) using ISOGEN with a spin column (Nippon Gene, Tokyo, Japan). Reverse transcription was performed using SuperScript III (Invitrogen, Waltham, Massachusetts, USA). PCR and quantification were performed using a StepOnePlus Real-Time PCR System (Applied BioSystems, Waltham, Massachusetts, USA) with THUNDERBIRD SYBR qPCR Mix (Toyobo, Osaka, Japan).

### Whole-mount in situ Hybridisation

Whole-mount *in situ* hybridisation was performed as previously described (Kawaguchi et al., 2008, 2005; Nagasawa et al., 2016). The three HCE cDNA had nucleotide sequences with >97% similarity to each other, therefore, we designed primers to amplify the common region (528bp) of AoHCE1, AoHCE2, and AoHCE3 (Table. S1) as an RNA synthesis template. RNA probe may cross-react with the mRNA of all AoHCEs. RNA probes were synthesised and labelled using the T7-ScribeTM Standard RNA IVT Kit (Madison, Wisconsin, United States) and Digoxigenin-11-UTP (Roche, Basel, Switzerland). Embryos at various stages were fixed in 4% paraformaldehyde overnight at 4°C, manually dechorionated, and immersed in 100% methanol at 4°C for one week. Proteinase K treatment was omitted to prevent unnecessary tissue degradation. Subsequently, embryos were hybridised at 55°C overnight in a hybridisation buffer (50% formamide, 5x SSC, 50 μg mL^-1^ heparin, 50 μg mL^-1^ tRNA, 0.1% Tween-20) with 0.5 ng μL^-1^ DIG-labelled RNA probe. Embryos were washed four times each with solution I (50% formamide, 2xSSC, 0.1% Tween-20), solution II (2xSSC, 0.1% Tween-20), and solution III (0.2xSSC, 0.1% Tween-20) at 68℃ for 30 minutes. After wash in PBST at room temperature, embryos were preabsorbed in 1% blocking reagent at room temperature for 90 minutes and immersed in anti-DIG antibody solution (using a 1:8000 dilution in PBST) at 4℃ for overnight.

After washing several times with PBST, the embryos were incubated in AP buffer (100 mM Tris ⁄ HCl pH 9.5, 50 mM MgCl2, 100 mM NaCl, and 0.1% Tween-20) for 10 minutes and stained with NBT ⁄ BCIP solution (1:50 in AP buffer).

### Fluorescent Labelling of Hatching Gland Cells

To visualise hatching gland cells in clown anemonefish embryos, we injected tol2 mRNA and an EGFP expression vector (pAoHCE2-EGFP-tol2; accession No.LC791262) or an EGFP/mCherry co-expression vector (pAoLCE-EGFP-AoHCE2-mch-tol2; accession No. LC794202) into fertilised eggs before the first cleavage at a concentration of 50 ng μL^-1^ for tol2 mRNA, and 5 ng μL^-1^ for pAoHCE2-EGFP-tol2 or pAoLCE-EGFP-AoHCE2-mch-tol2. These plasmids contained fluorescent protein genes driven by the AoLCE or AoHCE2 promoters and were integrated into the genomic DNA by tol2 transposase (Kawakami et al., 2004; Urasaki et al., 2006). Tol2 transposase mRNA was synthesised from pCS-TP (Kawakami et al., 2004) using an mMessage mMachine SP6 Kit (Ambion, Austin, Texas, USA). Fluorescence was observed using a fluorescence stereomicroscope (SZX16, OLYMPUS, Tokyo, Japan) with GFP (excitation: BP460-495, emission: BA510IF, OLYMPUS) and RFP (excitation: 562/40, emission: 641/75, OLYMPUS) filter sets.

### Chorion Digestion Analysis and N-terminal Amino Acid Sequences

Eggs were collected from the spawning plate in the late afternoon on the eighth day after fertilisation. Pre-hatching egg envelopes were isolated using fine tweezers. Post-artificial-hatching egg envelopes were obtained through the artificial hatching method (Yamanaka et al., 2021), whereas post-natural-hatching egg envelopes were collected from the spawning plate after larvae naturally hatched in the aquarium on the eighth night post spawning. Subsequently, the egg envelopes were treated with an SDS solution comprising 50 mM Tris-HCl, 2% SDS, 10% glycerol, and 0.005% bromophenol blue at 100°C for 5 minutes. SDS-PAGE was performed using a 5-20% gradient gel (E-T/R/D520L; ATTO, Tokyo, Japan). After electrophoresis, the gel was stained with Coomassie Brilliant Blue (CBB) to visualise the chorion digestion patterns. Additionally, to determine the N-terminal amino acid sequences, egg envelope proteins wered electroblotted onto a 0.45 μm PVDF membrane (Amersham, Princeton, NJ, UK) after SDS-PAGE. The membrane was then stained with CBB, and the band at approximately 30–40 kDa was excised and subjected to N-terminal amino acid sequencing using a SSPQ-20 protein sequencer (Shimadzu, Kyoto, Japan).

### Effect of Environmental Cues on Chorion Digestion

Eggs were collected from the spawning plate on the hatching day. Subsequently, eggs were divided into four batches and incubated for 90 minutes at 27°C under four conditions: (1) lights on without shaking, (2) lights off without shaking, (3) lights on with shaking, and (4) lights off with shaking. Lights-off conditions were achieved by completely wrapping the container with aluminium foil, whereas lights-on conditions approximated normal room brightness. Shaking was performed at 120 rpm using a Multi shaker MMS-110 (TOKYO RIKAKIKAI CO., LTD, Tokyo, Japan). After incubation, egg envelopes were isolated from the embryos using fine tweezers, and the extent of chorion digestion was assessed by SDS-PAGE as described above.

### Effect of Ca^2+^ on Chorion Digestion

Eggs collected from the spawning plate on the hatching day were placed in seawater containing either 2 μM ionomycin (in DMSO) or 0.1% (v/v) DMSO for 90 minutes under both light and dark conditions without shaking. After incubation, the extent of chorion digestion was assessed by SDS-PAGE.

### Effect of Mechanical Water Agitation on Chorion Digestion

Eggs collected from the spawning plate on the hatching day were subjected to shading for various durations (0–80 minutes), with or without shaking. After incubation, the extent of chorion digestion was assessed by SDS-PAGE.

## Result

### Phylogenetic Analysis of Hatching Enzymes

The hatching enzyme-like genes were identified in the clown anemonefish genome database through BLAST searches based on their similarity to medaka HCE and LCE. The phylogenetic tree revealed the presence of five distinct clades: HCE, LCE, patristacin, nephrosin, and pactacin (Fig. 1). Within the HCE clade, four hatching enzyme-like proteins (XP_023118123, XP_054868003, XP_023118132, and XP_054868280) were identified. XP_023118132 and XP_054868280 are variants transcribed from the same gene. In the LCE clade, a single hatching enzyme-like protein (XP_023133885) was found. These genes were named as *Amphiprion ocellaris* HCE (AoHCE) 1–3 and AoLCE. The remaining seven hatching enzyme-like proteins were distributed among nephrosin and pactacin, which belonged to the c6 astacin family (Fig. 1).

**Fig. 1.**
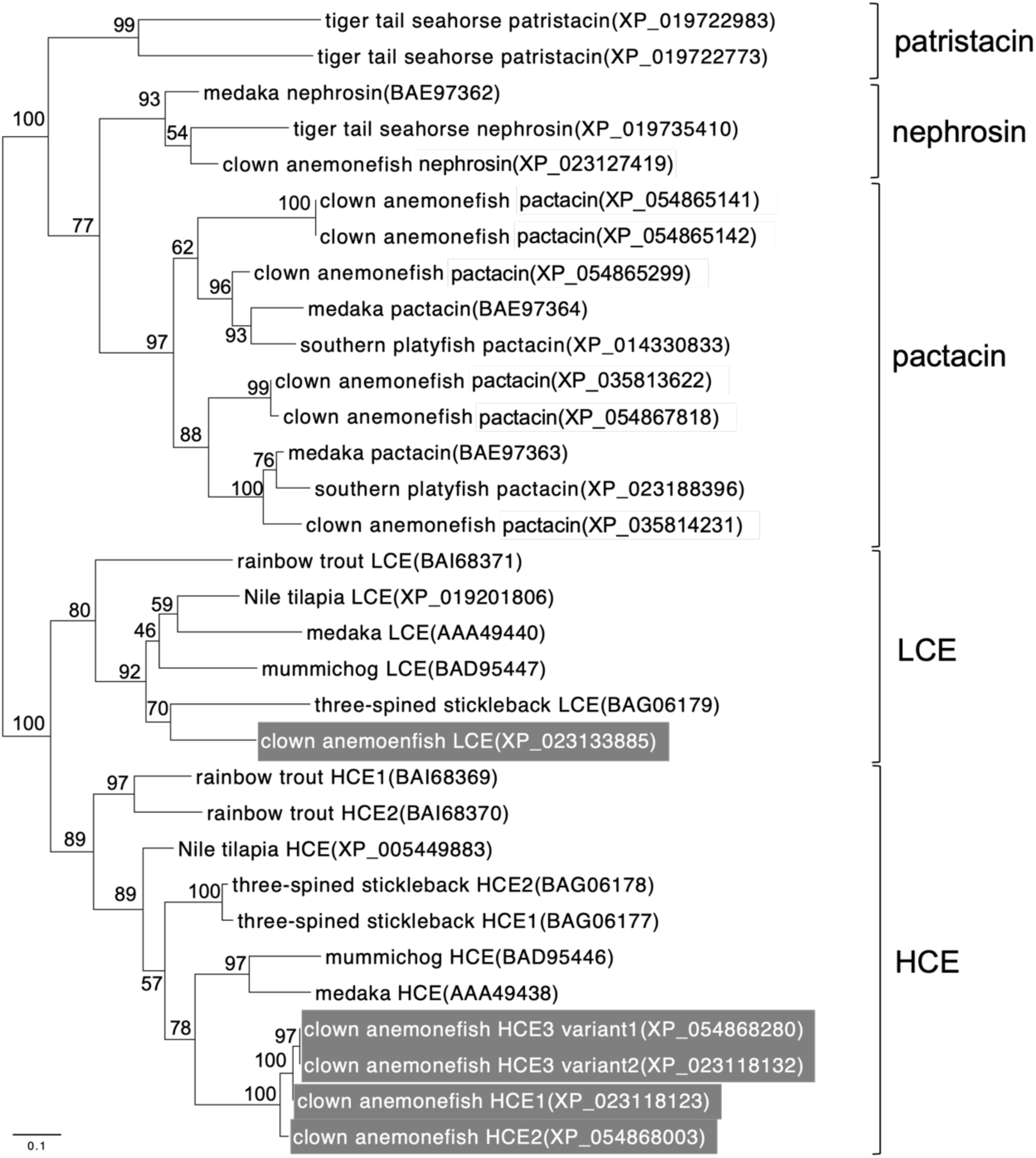
A phylogenetic tree of C6AST and hatching enzyme genes. The phylogenetic tree was constructed from amino acid sequences of hatching enzyme-like genes of clown anemonefish and C6ASTs of other teleost fishes by the maximum likelihood method. HCEs and LCE of clown anemonefish were shaded in dark grey.

### Temporal Expression Patterns of Hatching Enzymes

The expression profiles of HCE1, HCE2, HCE3, and LCE showed an upward trend during embryonic development from 1 to 2 dpf, followed by a gradual decline until the hatching day (8 dpf) (Fig. 2). At 1 dpf, several somites and eye primordia were observed. At 2 dpf, the beginning of black pigmentation on yolk were observed. The expression levels of these four enzymes were low post-hatching (9–10 dpf) (Fig. 2).

**Fig. 2.**
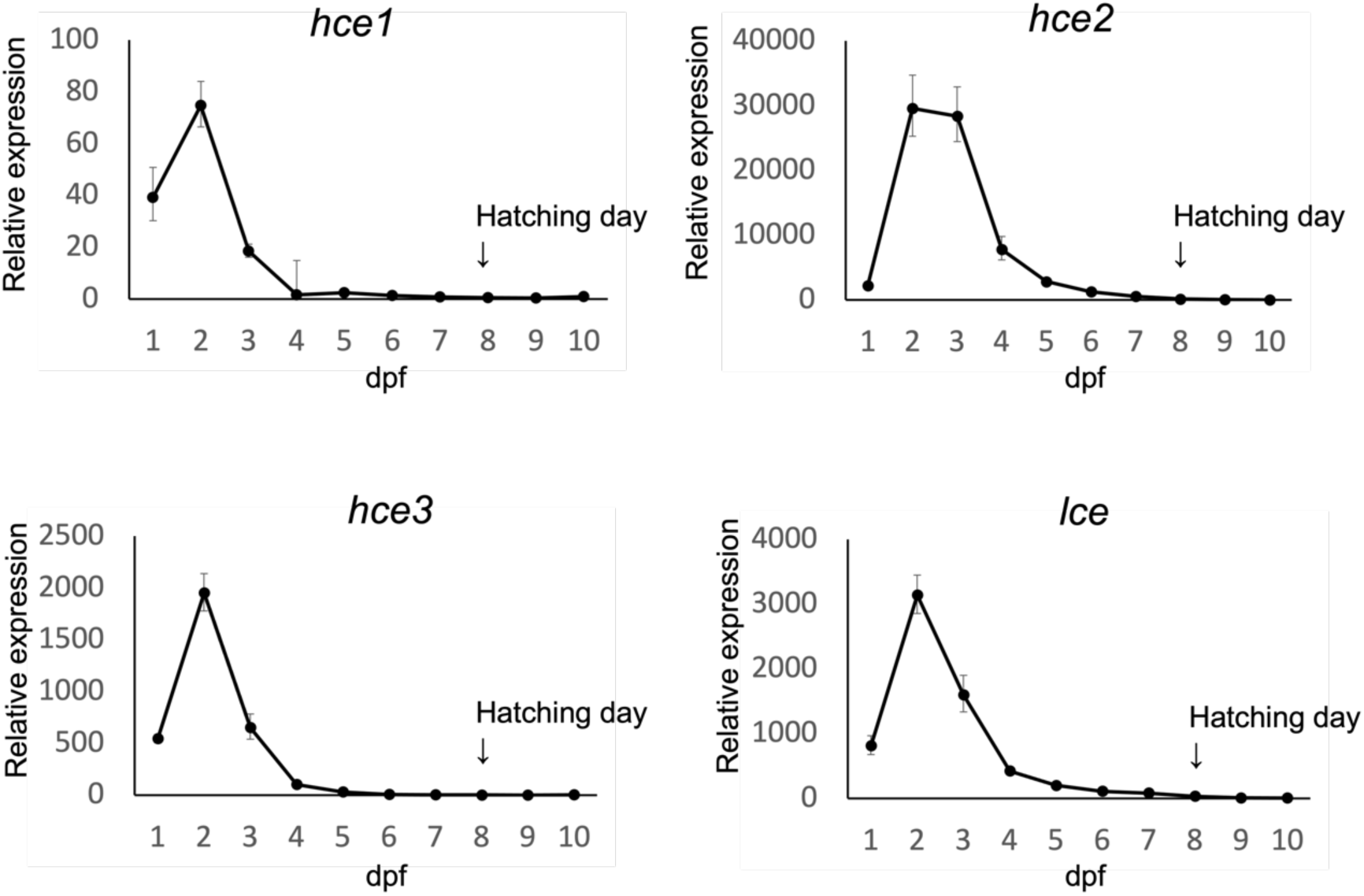
The temporal expression pattern of hatching enzymes in clown anemonefish. The relative expression level was determined by using the delta-delta Ct method. Data are presented as fold changes (2^(-ΔΔCt)) relative to 10 dpf (days post fertilisation) samples, with error bars representing standard deviations (n=3). Beta-actin2 (actb2) was used as an internal control. The expression of all hatching enzyme genes peaked on 2 dpf (when somite segmentation is completed and black pigmentation on yolk begins) and declined towards hatching day (8 dpf).

### Localisation of Hatching Gland Cells

Fluorescence labelling driven by AoHCE2 promoters and *in situ* hybridisation using an anti-sense probe for AoHCEs revealed that the hatching gland cells were primarily localised on the tail, extending from the upper part of the anus to the edge of the caudal fin on the ventral side (Fig. 3A–D). EGFP driven by the AoLCE promoter and mCherry driven by the AoHCE2 promoter were co-expressed in these cells (Fig. 3D-4–6). No signal was observed with the sense probe (data not shown).

**Fig. 3.**
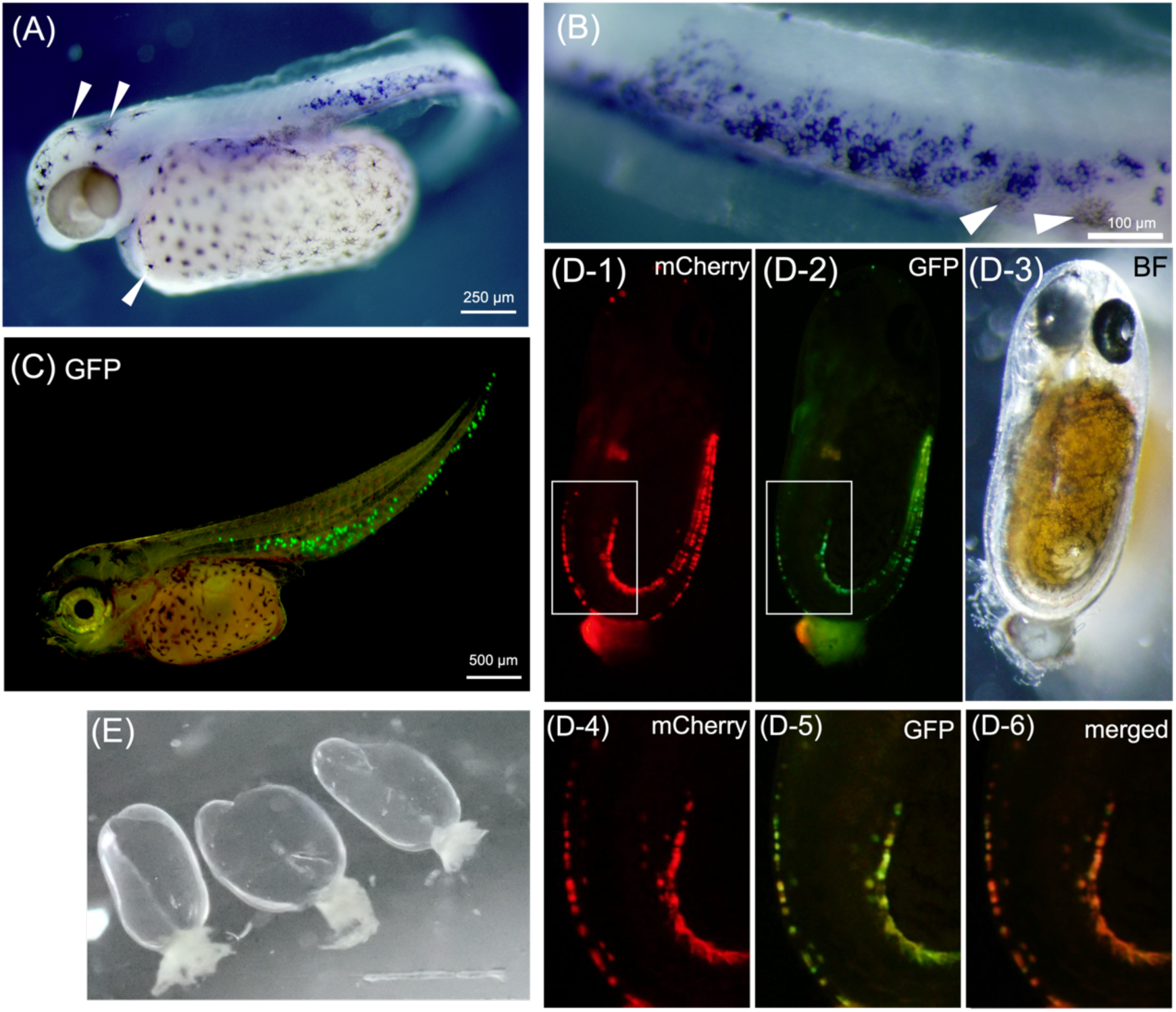
The localization of hatching gland cells and egg envelopes after hatching. Embryos at 3 dpf (A), 6 dpf (B,C,D). Hatching enzyme expressions (blue signals) were detected by whole-mount *in situ* hybridisation using antisense RNA probe for HCE (A,B). *Arrowhead*: black pigment. C: Hatching gland cells labelled by EGFP driven by HCE2 promoter. D: Hatching gland cells co-labelled by EGFP driven by LCE promoter and mCherry driven by HCE2 promoter. These two hatching enzyme genes are expressed in the same cell (D-4,5,6). E: The egg envelopes after hatch. They were consistently broken along the midpoint of the ellipsoid’s long axis.

**Fig. 4.**
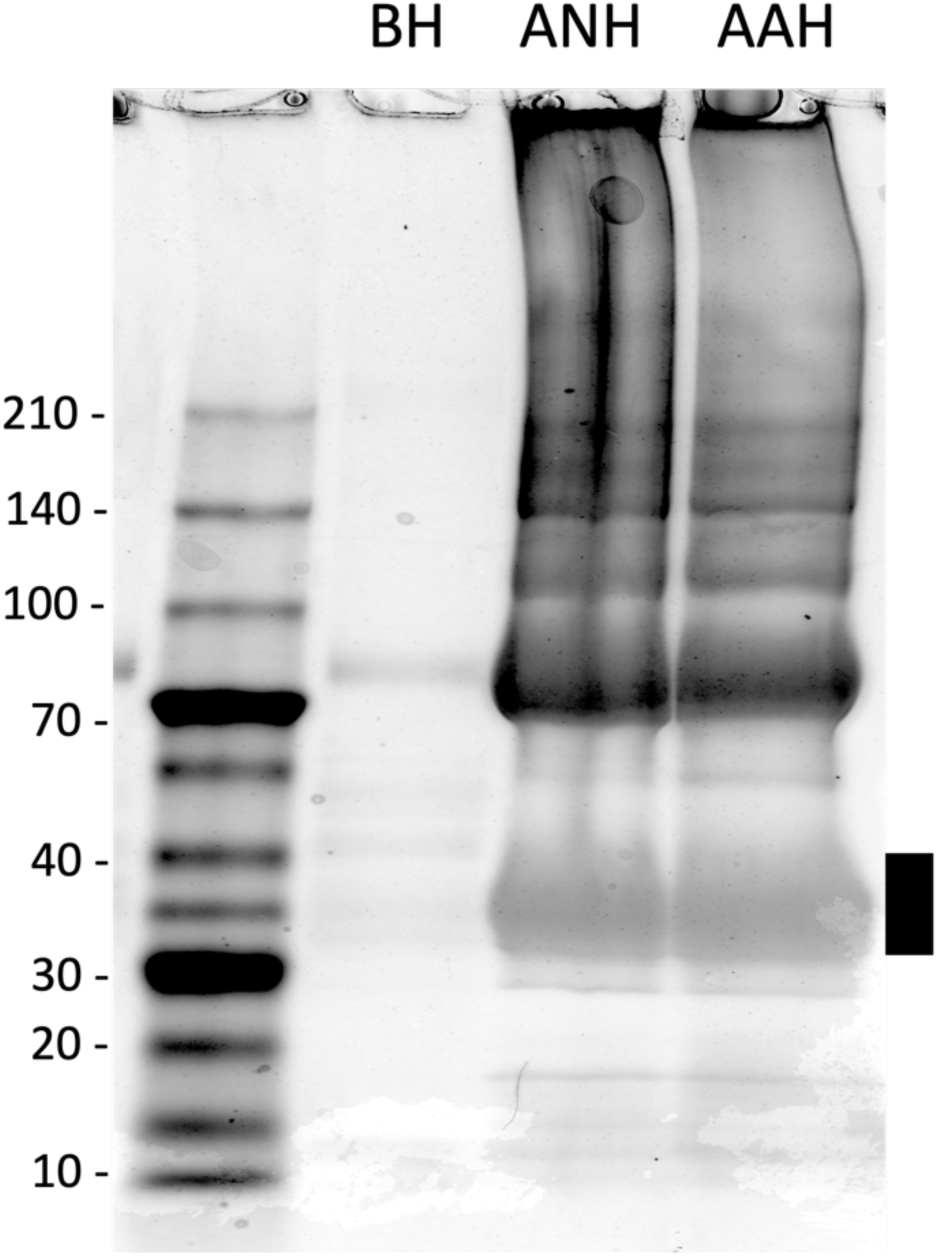
SDS-PAGE patterns of proteins in egg envelopes before hatch (BH), after natural hatch (ANH), and after artificial hatch (AAH). Egg envelopes after both natural and artificial hatch showed several bands but egg envelopes before hatch did not. The black line indicates the 30-40 kDa bands used for analyzing of N-terminal amino acid sequence (Table. 1.).

### Analysis of Digested Proteins of Chorion

Significant differences were observed in band patterns between egg envelopes before and after hatching. Post-hatching egg envelopes (ANH and AAH in Fig. 4) exhibited multiple bands, whereas egg envelopes before hatching (BH in Fig. 4) displayed only a few bands. According to the protein sequencing result, the band at approximately 30–40 kDa showed multiple amino acid sequences. Subsequently, we predicted all possible combinations of amino acids at each locus and conducted a BLAST search using these sequences as queries (Fig. S1). Three N-terminal amino acid sequences were predicted as the fragment of AoChgH (XP_023152162), AoChgHm (XP_023147961), and AoChgL (XP_023147962) (Table 1). The molecular weights from the predicted amino terminals to the C-terminals of each Chg proteins can be estimated to be 39.7, 42.0, 42.1 kDa, respectively, which are close to the molecular weights estimated by SDS-PAGE. Additionally, these N-terminal sequences were consistent with the cleavage sites of HCE on each Chg protein in mummichog (Fig .S2) (Kawaguchi et al., 2010b).

**Table. 1.** Predicted cleavage site and predicted molecular weight of each Chg digested products based on the N-terminal amino acid sequences around 30-40 kDa bands. An overlap of amino acid sequences was detected around 30-40 kDa band in Fig. 4, which were predicted to contain partial sequences of AoChgH, AoChgHm, and AoChgL. The predicted cleavage sites from N-terminal sequences are consistent with those of mummichog (Fig. S2). X: Residues could not be detected by protein sequencer.

| Predicted N-terminal sequences | Chg genes | site | predicted MW<br>(kDa) |
| --- | --- | --- | --- |
| HPALRNxDVDLSV | AoChgH(XP_023152162) | Q205/H206 | 39.7 |
| EPQKPQPxxDAFH | AoChgHm(XP_023147961) | K84/E85 | 42.0 |
| EPPxQKQTFEQPL | AoChgL(XP_023147962) | Y43/E44 | 42.1 |

### Effect of Darkness on Chorion Digestion

Hatching occurred when eggs were shaken and shaded, while other conditions did not induce hatching, which is consistent with the previous study (Yamanaka et al., 2021). SDS-PAGE analysis of egg envelopes revealed distinct band patterns under the different experimental conditions (Fig. 5A). Under dark condition, multiple bands were detected, regardless of shaking (Fig. 5A), resembling the pattern observed in egg envelopes after hatching (Fig. 4). Conversely, limited and faint band patterns were observed under light conditions (Fig .5A).

**Fig. 5.**
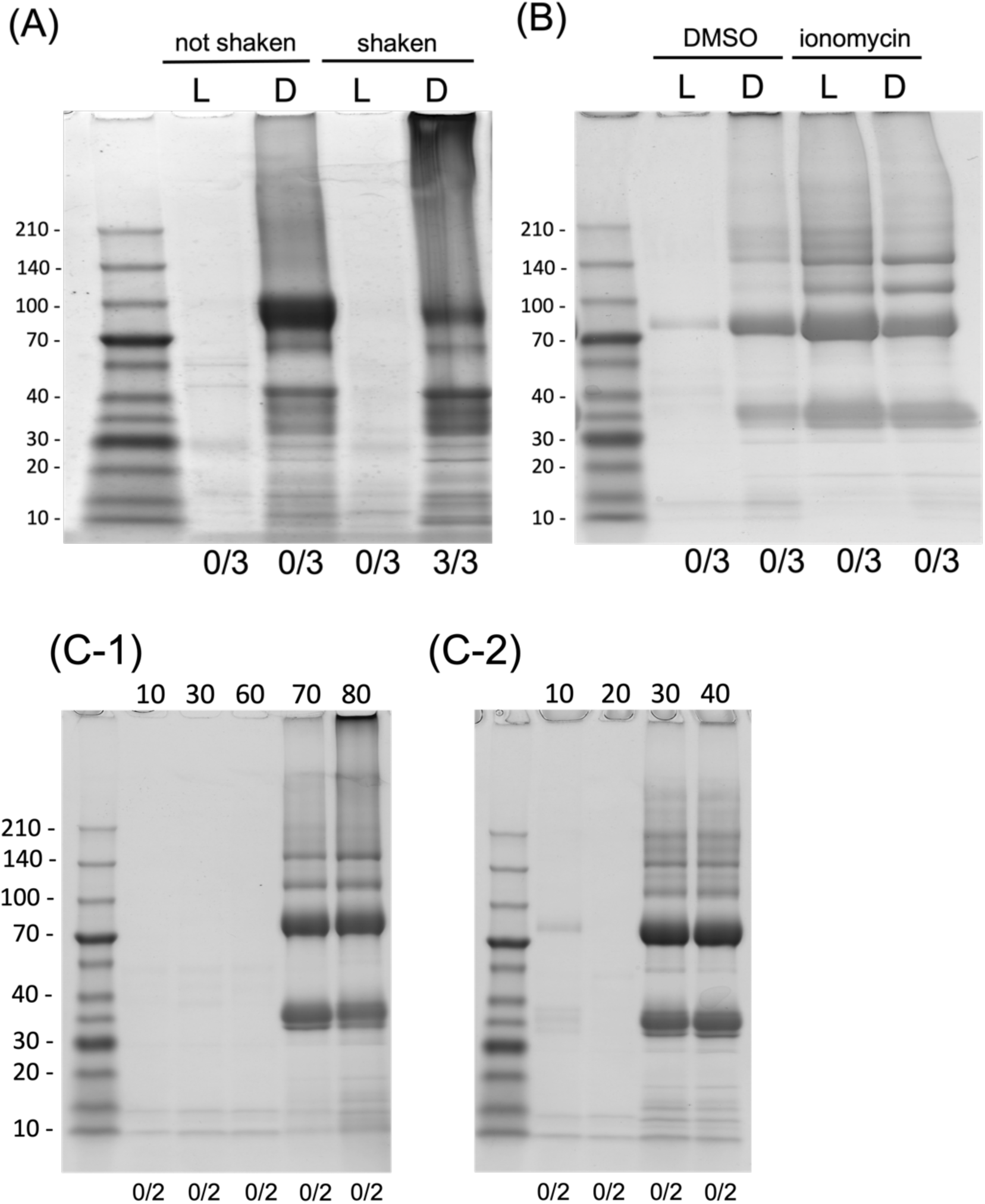
Chorion digestion patterns in various conditions. The numbers under each lane indicate the hatching rate (hatched embryos/total embryos). (A) The chorion digestion patterns in response to shading and shaking manipulation. L: Lights-on condition. D: Drak condition. Dark conditions induced chorion digestion with or without shaking treatments, whereas light condition did not. Hatching occurred only when eggs were subjected to shading and shaking. (B) The chorion digestion patterns in response to ionomycin. Ionomycin triggered the chorion digestion regardless of light conditions, but DMSO (solvent) did not. (C) The time series (minutes) of chorion digestion patterns under light off condition without shaking treatment (C-1) and with shaking treatment (C-2). Chorion digestion occurred earlier with shaking treatment compared to without shaking treatment.

### Effect of Ca^2+^ on Secretion of Hatching Enzyme

In the presence of DMSO, chorion digestion occurred only in the dark. Ionomycin, which facilitates Ca^2+^ transport into cells (Liu and Hermann, 1978) and induces hatching enzyme secretion (Iuchi et al., 1985), triggered chorion digestion irrespective of light conditions (Fig. 5B).

### Effect of Mechanical Agitation on Chorion Digestion Activity

Analysis of the SDS-PAGE results revealed distinct outcomes based on the presence or absence of mechanical agitation. When the eggs were shaded but not shaken, band patterns characteristic of chorion digestion were observed after 70 minutes of treatment (Fig. 5C-1). In contrast, when eggs were subjected to both shading and shaking, bands were detected after only 30 minutes of treatment (Fig. 5C-2), indicating mechanical water agitation expedites chorion digestion.

## Discussion

### Expression Pattern of Hatching Enzyme

Phylogenetic analysis based on amino acid sequences revealed three genes clustered within the HCE clade, whereas a single gene was assigned to the LCE clade (Fig. 1). These genes were identified as AoHCE1 (XP_023118123), AoHCE2 (XP_054868003), AoHCE3 (XP_023118132, XP_054868280), and AoLCE (XP_023133885). All hatching enzyme genes contain consensus motifs, including HExxHxxGFxHExxRxDR, SxMHY, and six cysteine residues that are characteristic of c6 astacin genes, including hatching enzymes (Fig. S3). Three HCE genes displayed an intron-less structure, whereas the LCE gene featured an 8-exon-7-intron structure. These gene structures were consistent with the phylogenetic-specific gene structure of hatching enzymes observed in euteleostei (Kawaguchi et al., 2010a; Sano et al., 2014). Expression analysis revealed that these four hatching enzymes exhibited high mRNA levels in the early developmental stages (1–2 dpf), which gradually decreased as hatching day approached (Fig. 2). To further investigate the localisation of the hatching gland cells which express hatching enzymes, we conducted whole-mount *in situ* hybridisation using an anti-sense RNA probe for AoHCEs. Our results demonstrated that hatching gland cells were unicellular cells arranged along the ventral side of the lateral line, extending from the anterior of the anus to the tip of the tail (Fig. 3A,B). This localisation pattern was further corroborated by fluorescence labelling with EGFP driven by the AoHCE2 promoter (Fig. 3C). Additionally, our observations showed co-labelling of hatching gland cells with EGFP and mCherry driven by the AoLCE and AoHCE2 promoters, respectively (Fig. 3D), indicating the coincidental expression of all four hatching enzymes. The location of hatching gland cells varies significantly across fish species (Inohaya et al., 1999, 1997, 1995; Kawaguchi et al., 2009, 2005; Martin et al., 2011) and may be closely related to factors such as egg shape, chorion thickness, manner of chorion breakage, and nature of hatching enzyme activity (Korwin-Kossakowski, 2012). In clown anemonefish, embryonic development involves a folded tail within an ellipsoidal egg capsule. Larvae emerge tail-first (Salis et al., 2021), and egg envelopes are consistently broken along the midpoint of the ellipsoid’s long axis (Fig. 3E). These observations suggest that hatching gland cells located on the tail possibly play a pivotal role in chorion digestion around the tail region and contribute to chorion rupture by stretching the tail to facilitates rapid hatching in response to environmental cues.

### Biochemical Properties of Hatching Enzyme

In fish, the egg envelope proteins (ChgH,ChgHm, and ChgL) contain zona pellucida (ZP) domain. ChgH and ChgHm belong to the ZPB subfamily, whereas ChgL belongs to the ZPC subfamily. Following fertilisation, these protein subunits rapidly cross-link with each other to shield the embryo from mechanical and chemical stresses (Hamazaki et al., 1987; Murata et al., 1997, 1995; Sugiyama et al., 1998). Studies in medaka and mummichog have shown that intact egg envelopes are insoluble in SDS, however, these can be solubilised through the coordinated digestion of two hatching enzymes, HCE and LCE(Kawaguchi et al., 2010b; Yasumasu et al., 2010). In clown anemonefish, a notable disparity in band patterns was observed in the lysate from the egg envelope before and after hatching (Fig. 4). The post-hatching egg envelopes exhibited several distinct bands, whereas those before hatching exhibited only a few bands. Multiple amino acid sequences were predicted at approximately 30–40 kDa bands, which matched the partial sequences of AoChgH, AoChgHm, and AoChgL (Table 1, Fig. S1). Aligning these sequences with the choriogenin amino acid sequences of mummichog revealed that these N-terminal sequences corresponded to the conserved cleavage site targeted by HCE in mummichog and medaka (Kawaguchi et al., 2010b) (Fig. S2). This suggests that the cleavage site recognized by HCE is conserved among clown anemonefish, mummichog, and medaka. Furthermore, we examined the biochemical properties of the hatching enzymes of clown anemonefish using inhibitors. Previous studies have shown that hatching enzymes are zinc metalloproteases, and that activity can be suppressed by chelating agents such as EDTA (Martin et al., 2011; Yamagami, 1973; Yasumasu et al., 1989a, 1989b). Additionally, hatching enzymes exhibit varying degrees of salt dependence across fish species, adapting to the specific salinity conditions of their respective environments (Kawaguchi et al., 2013). Our results demonstrated that chorion digestion activity was inhibited under low salinity conditions or in the presence of EDTA (Fig. S4). Consequently, the biochemical characteristics of hatching enzymes in clown anemonefish resemble those observed in other fish species.

### Effects of Environmental Cues on Hatching Processes

Some fish species employ hatching strategies in which environmental cues play an essential role. The hatching of clown anemonefish is also affected by environmental cues, including complete darkness and water agitation (Fobert et al., 2019; Yamanaka et al., 2021). However, the precise physiological mechanisms underlying how environmental cues affect hatching processes remained unclear. As described above, we confirmed similarities in the characteristics of hatching enzymes and the chorion digestion mechanism in clown anemonefish with other teleost fish species.

Therefore, we focused on roles of environmental cues in the secretion of hatching enzymes and the chorion digestion. We performed SDS-PAGE to examine the extent of chorion digestion after several conditioned incubations. Interestingly, the dark conditions induced chorion digestion irrespective of water agitation, whereas the lights-on condition did not induce chorion digestion (Fig. 5A). This intriguing result suggests that darkness triggers the secretion of hatching enzymes or enhances the biochemical activity of hatching enzymes. Given the absence of a specialised domain in hatching enzymes of clown anemonefish, it is improbable that the biochemical activity of hatching enzymes is directly affected by light conditions. Therefore, we investigated the relationship between dark stimuli and the secretion of hatching enzymes. Ca^2+^ levels have been implicated in the secretion of hatching enzymes in several fish species (Iuchi et al., 1985; Schoots et al., 1981). Calcium ionophore, a potent inducer of hatching enzyme secretion in other fish species, also resulted in chorion digestion independently of light conditions in clown anemonefish (Fig. 5B). Additionally, hatching enzymes are released from hatching gland cells as mature enzymes and promptly initiate the digestion of chorion in medaka (Iuchi et al., 1982). Therefore, it is suggested that hatching enzymes are stored within hatching gland cells and is released following dark stimulation via a signalling mechanism associated with increased Ca^2+^ level. The elevation of Ca^2+^ level may plays a critical role in the transmission of dark stimuli and/or in the secretory processes of hatching gland cells. Subsequently, we investigated the effect of water agitation on hatching processes. We compared the time required for chorion digestion between eggs subjected to water shaking and those without shaking under lights-off conditions. Remarkably, shaking treatment under dark conditions expedited chorion digestion (Fig. 5C), indicating that water agitation enhances chorion digestion activity by assisting enzyme secretion and/or enzymatic activity. In summary, our findings demonstrate that darkness triggers hatching enzyme secretion, whereas water agitation expedites chorion digestion. Both these factors contribute significantly to chorion digestion, functioning as essential environmental cues for hatching in clown anemonefish. Notably, hatching occurred efficiently regardless of embryo density (Fig. S5), indicating that each embryo independently senses environmental cues and tightly regulates its response, however, they are not affected by the hatching success of other embryos. The stringent regulation of physiological processes by environmental cues underscores the rapid and synchronised hatching strategy employed by clown anemonefish.

### A New Model for Studying ECH

ECH has been the subject of significant investigation in various fish species, with studies mainly focusing on its behavioural and ecological implications(Dimichele and Taylor, 1980; Griem and Martin, 2000; Martin et al., 2011; Warkentin, 2011a, 2011b). Grunions, for instance, hatch immediately upon immersion in agitated seawater(Griem and Martin, 2000; Martin et al., 2011; Smyder and Martin, 2002), suggesting that ECH in this species predominantly affects a behavioural control mechanism rather than hatching enzyme secretion. Conversely, in mummichog, hypoxia serves as an environmental cue that triggers respiratory movement and the subsequent secretion of hatching enzymes (Dimichele and Taylor, 1981, 1980). However, it remained unclear whether mechanical stimulation directly promotes the secretion of hatching enzymes or whether more complex physiological processes, including neural pathways, are involved. Several fish species, such as halibut (Helvik and Walther, 1993, 1992), tropical pomacentrid (McAlary and McFarland, 1993), and spotted rose snapper (Duncan et al., 2008), have been reported to regulate hatching in response to light conditions, similar to the phenomena observed in clown anemonefish. Nevertheless, owing to the challenges in obtaining a constant supply of eggs and applying reverse genetic techniques, the molecular basis of the relationship between light conditions (or other environmental cues) and hatching remained elusive. On the other hand, in medaka and zebrafish, well-established model fishes in developmental and molecular biology, hatching timing can exhibit substantial variations of several hours or even days between individuals, suggesting relatively weaker regulation by environmental cues. In the present study, we demonstrated that the photo-environment plays a crucial role in regulating the secretion of hatching enzymes in clown anemonefish. These light-responsive behaviours likely involve intricate physiological processes associated with the perception of darkness by photoreceptors and subsequent signal transduction mechanisms such as hormonal and neural pathways. To elucidate the molecular basis of ECH, certain prerequisites, including the availability of eggs, whole-genome information, and the application of reverse genetics, are imperative. Clown anemonefish, notable for their ease of rearing and the frequency of breeding pairs spawning every two weeks, have garnered significant attention in the fields of developmental biology, genetics, and evolution. In recent years, complete sequencing of the clown anemonefish genome has been performed (Ryu et al., 2022; Salamin et al., 2023), and reverse genetic techniques, such as genome editing, have been successfully applied (Mitchell et al., 2021, 2023; Yamanaka et al., 2021). Therefore, the strong regulation of physiological processes by environmental cues, coupled with their suitability for molecular biological investigations, establishes clown anemonefish as a suitable model for unravelling the molecular basis of ECH.

## Acknowledges

We would like to express our thanks to Dr. S. Ansai for valuable research advice and supports.

## Competence interests

No competing interests declared.

## Author contributions

S.Yamanaka and M.Kinoshita designed the study and wrote the paper. S.Yamanaka., M.Kawaguchi, S.Yasumasu, and K.S. performed the experiments and analyzed the data.

## Funding

This work was supported by JSPS KAKENHI Grant Number 23KJ1220 (SY) and 22H02678 (MK).

## Supplemental protocols, table, and figures

### Effect of Salinity or EDTA on Hatching

Eggs collected from the spawning plate on the hatching day were shaken and shaded for 60 minutes in seawater at various salinity (0.6-2.4%) or EDTA concentrations (0-5 mM). Then, the hatching rates were recorded, and the extent of chorion digestion was assessed by SDS-PAGE.

### Effect of Egg Density on Hatching

Eggs collected from the spawning plate on the hatching day were shaken and shaded for 60 minutes with various densities (1–30 eggs in each container). Then, the hatching rates were recorded.

**Supplemental Table. S1.**
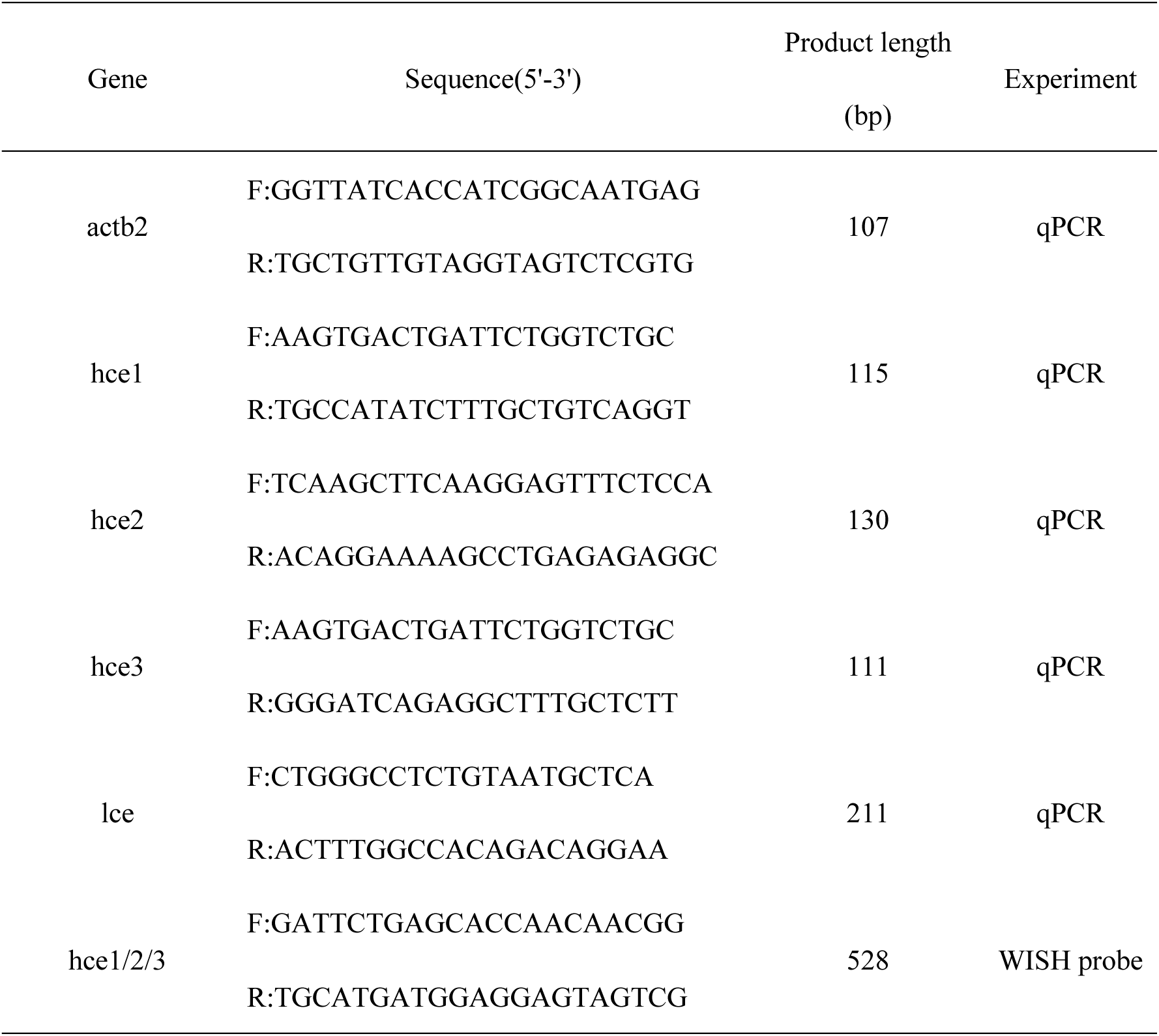
The primers used in RT-qPCR or RNA probe synthesize for whole-mount *in situ* hybridisation.

**Supplemental Fig. S1.**
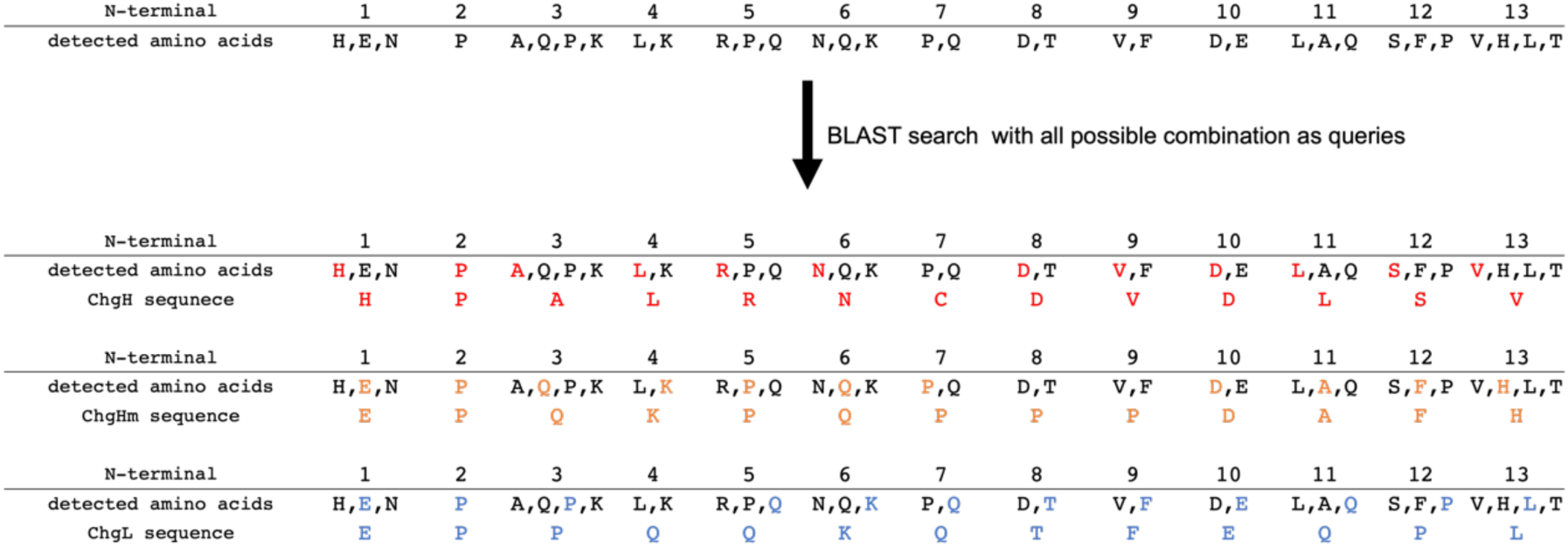
Prediction of amino acid sequences from around 30-40 kDa band. Multiple amino acids were detected in each locus from around 30-40 kDa band by protein sequencer indicating overlapping of several sequences. We conducted a BLAST search with all possible combination of amino acids as queries. As a result, it is strongly suggested that the obtained amino acid sequences contain the partial sequences of the three Chg proteins.

**Supplemental Fig. S2.**
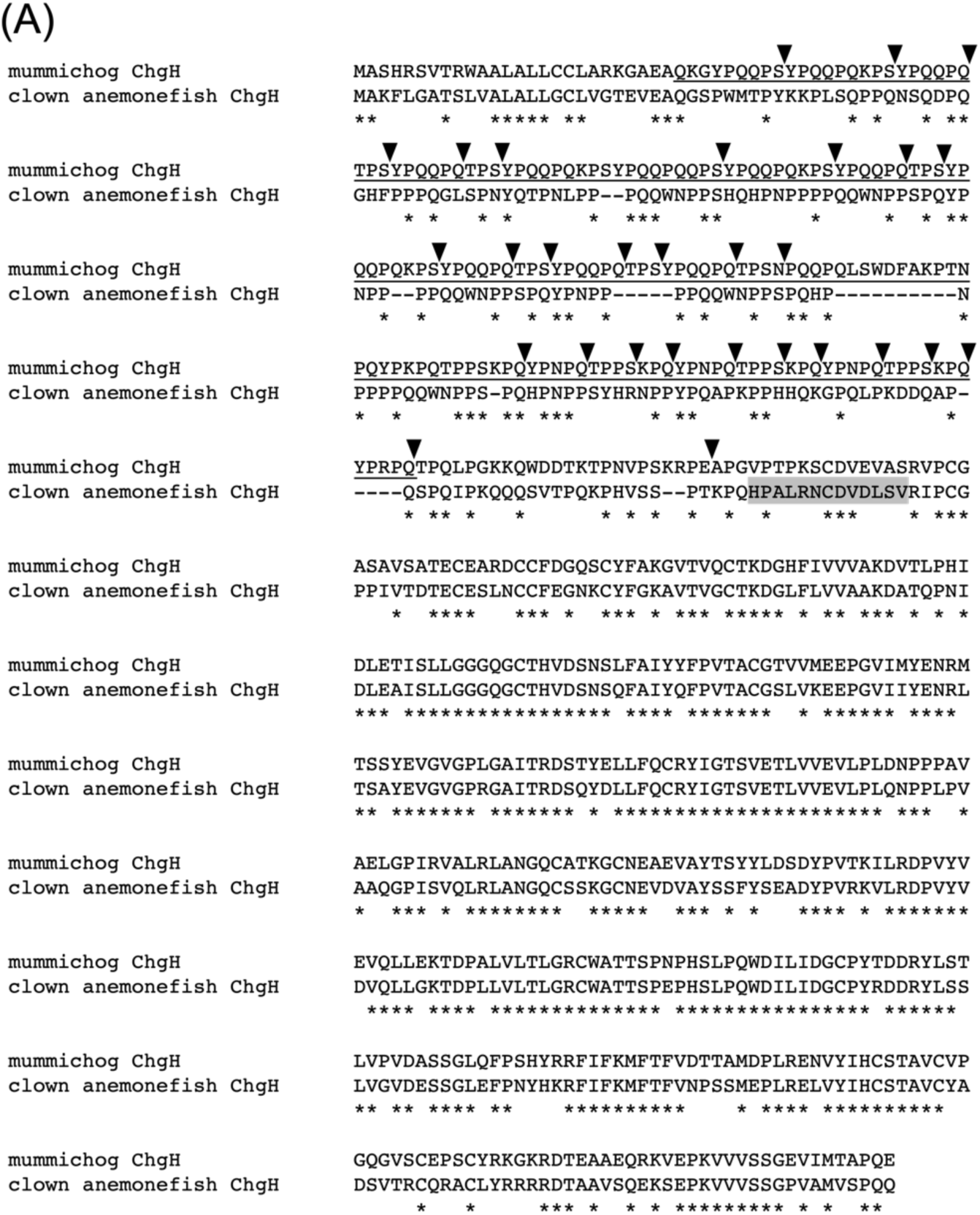
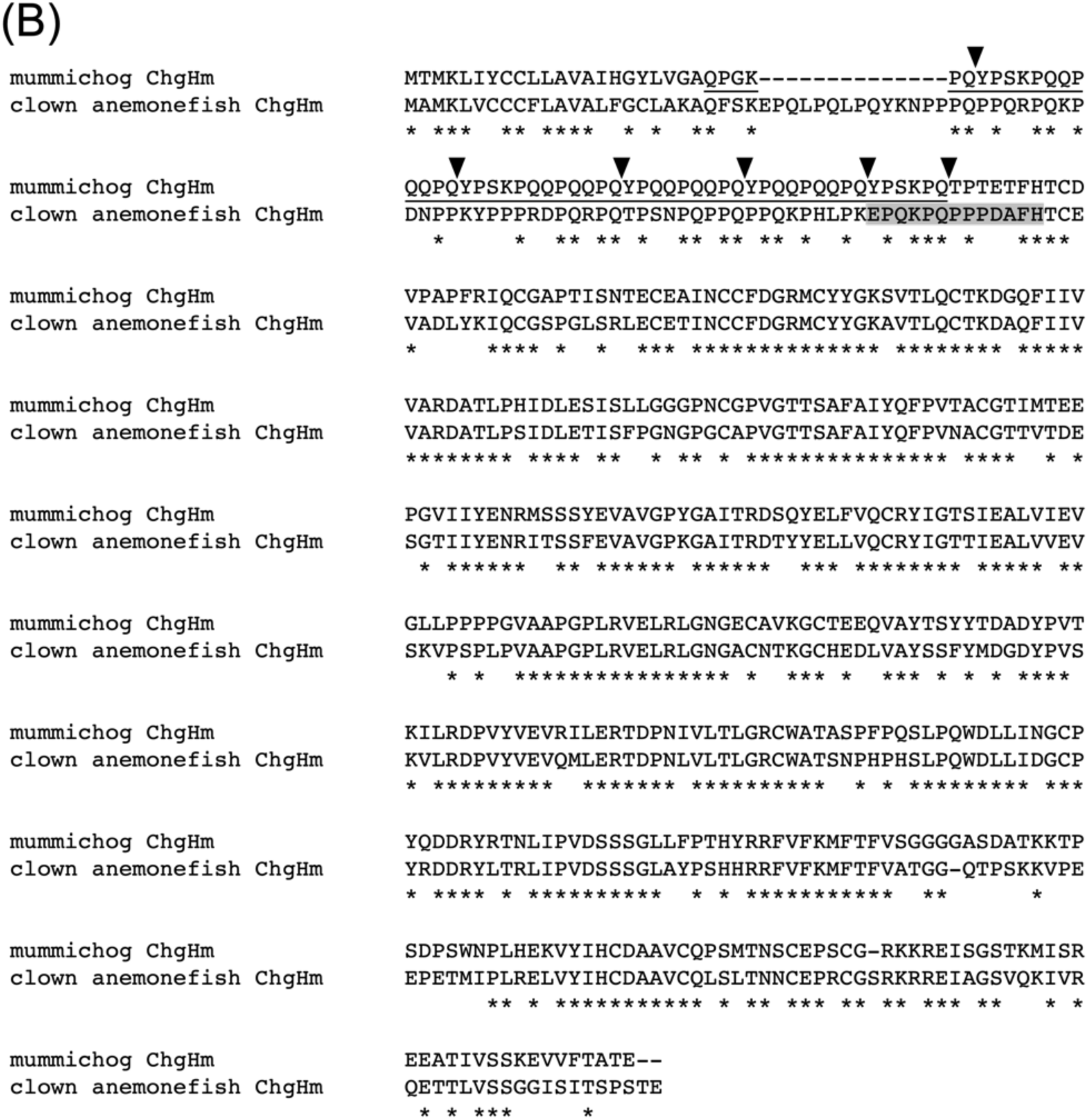
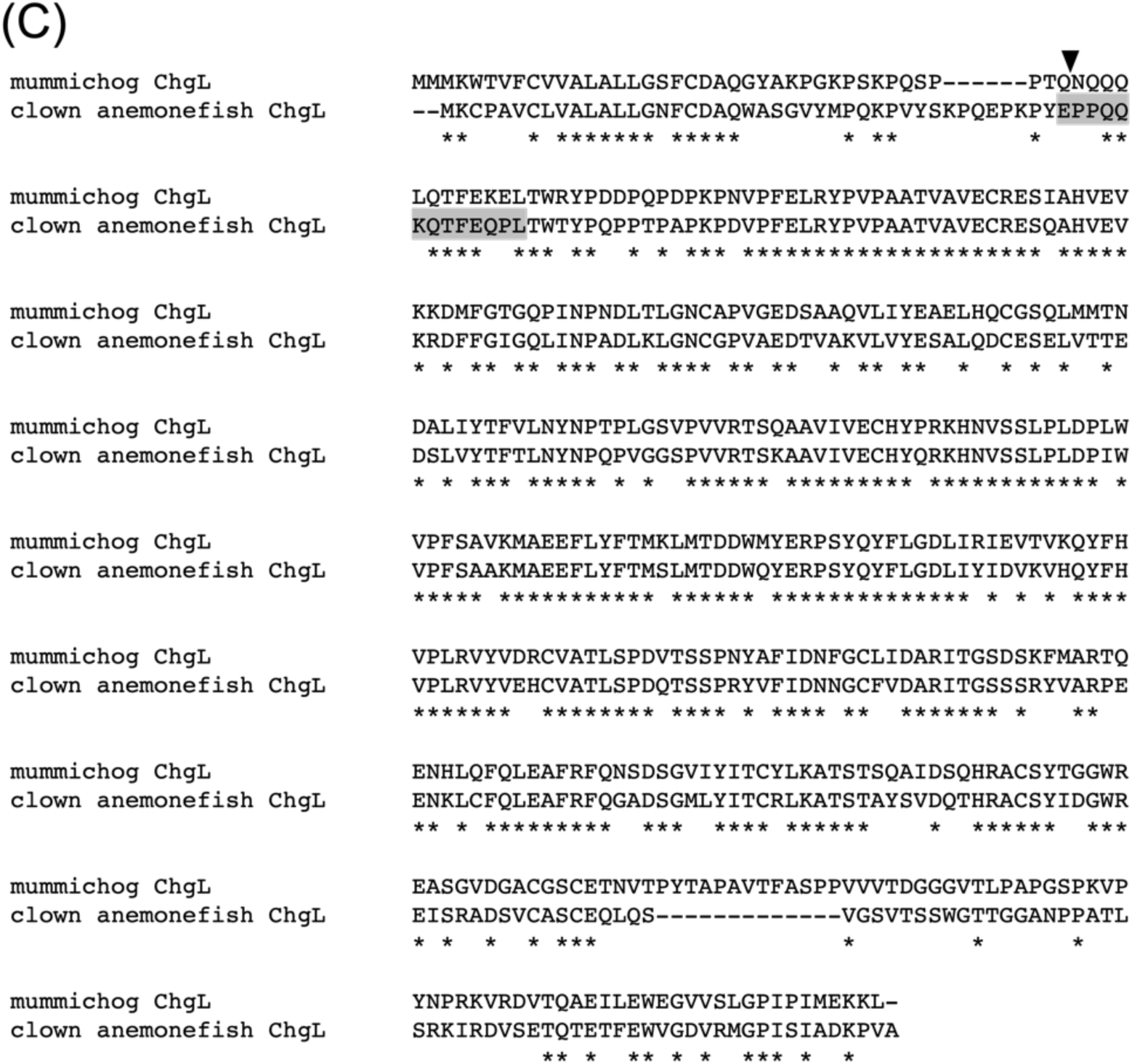
Alignment of amino acid sequences of ChgH (A), ChgHm (B), and ChgL (C) of mummichog and clown anemonefish. N-terminal sequences predicted in this study were shaded in dark gray. The cleavage sites by HCE in mummichog(Kawaguchi et al., 2010b) were presented as black arrowhead. Pro-X-Y repeat regions characteristic of ChgH and ChgHm were indicated by black lines.

**Supplemental Fig. S3.**
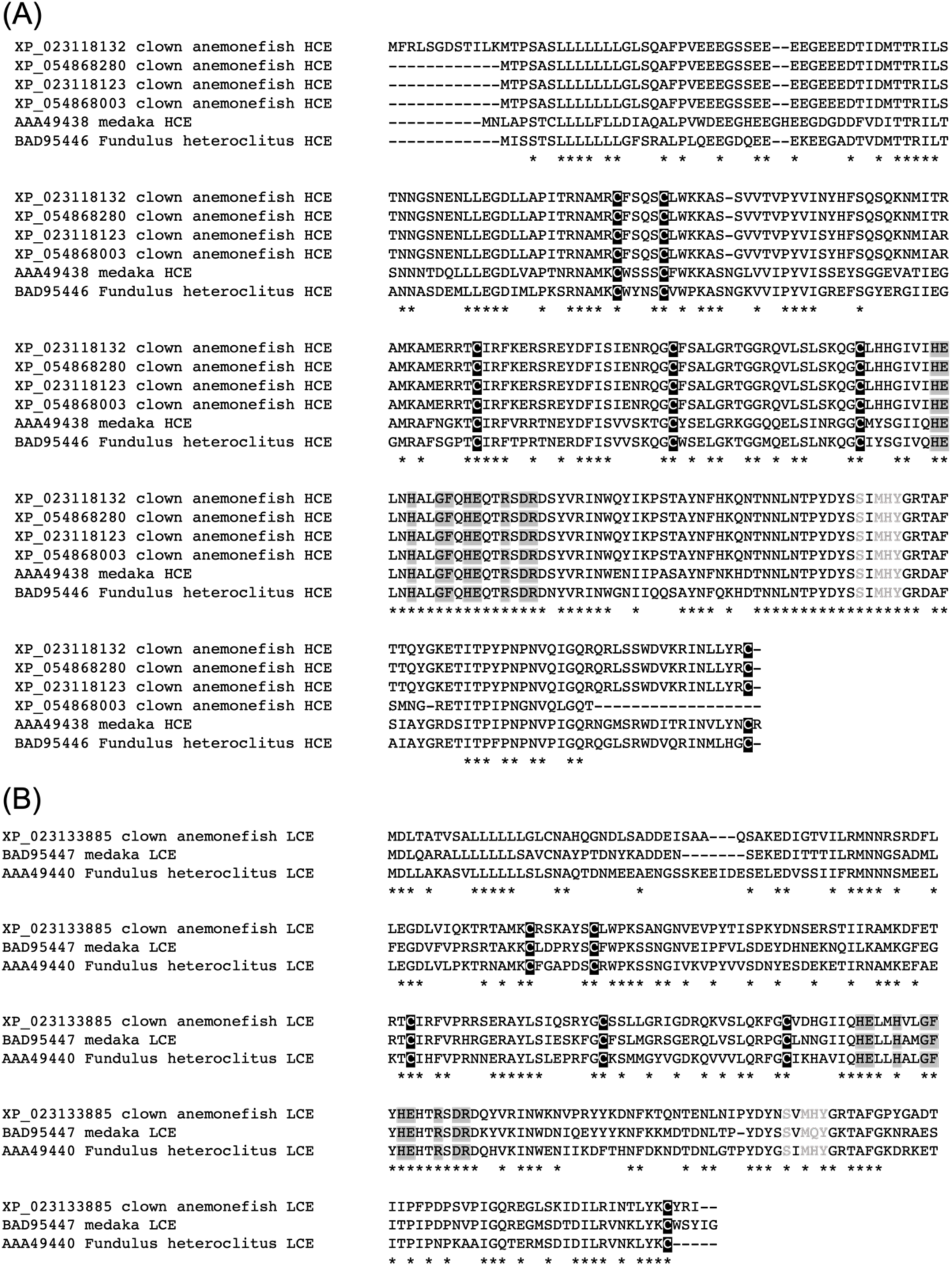
Alignment of amino acid sequences deduced from HCE and LCE cDNA of medaka, mummichog (*Fundulus heteroclitus*), and clown anemonefish. Consensus sequences of metal binding site HExxHxxGFxHExxRxDR and methionine turn SxMHY are shaded in dark gray and light gray, respectively. Cysteine residues conserved among c6ast genes are shaded in black.

**Supplemental Fig. S4.**
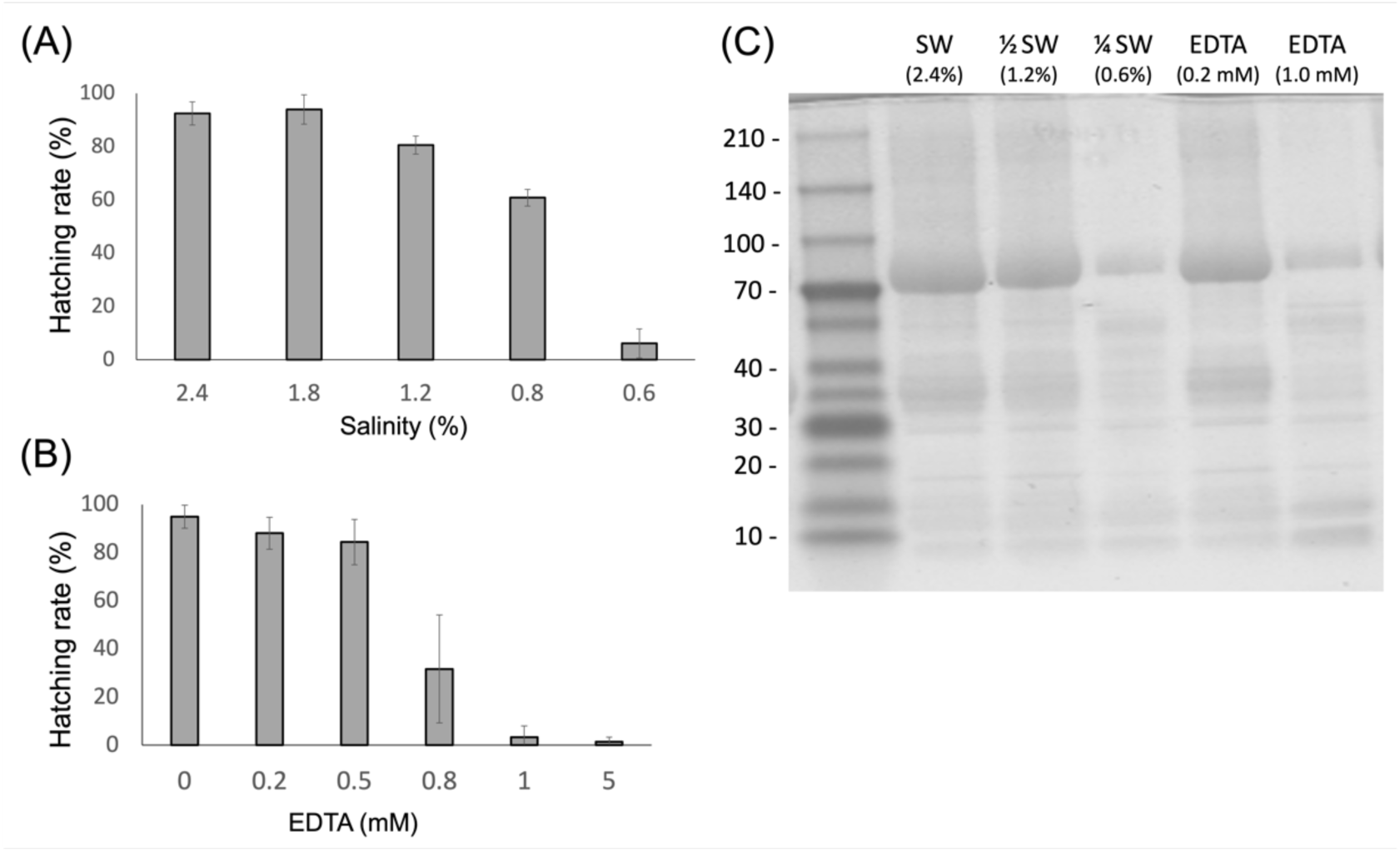
The effect of salinity and EDTA on hatching success and chorion digestion. A,B: The hatching rates under various salinity or EDTA concentrations. Hatching success decreased with decreasing salt concentration or increasing EDTA concentration. C: The chorion digestion patterns under various salinity or EDTA concentrations. Chorion digestion activity decreased with decreasing salt concentration or increasing EDTA concentration.

**Supplemental Fig. S5.**
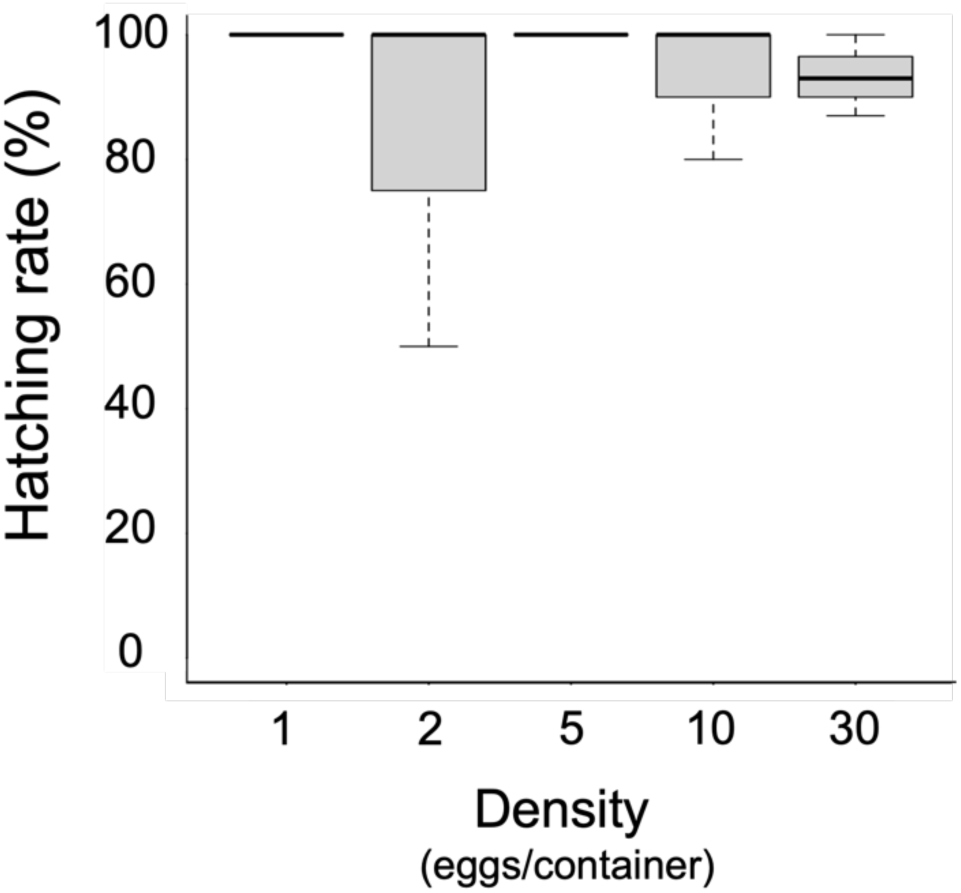
Hatching rates at different egg densities per container. Hatching occurred independently of egg density under light-off conditions with shaking treatment.

## Notes

### Competing Interest Statement

The authors have declared no competing interest.

